# Developing a Fully-glycosylated Full-length SARS-CoV-2 Spike Protein Model in a Viral Membrane

**DOI:** 10.1101/2020.05.20.103325

**Authors:** Hyeonuk Woo, Sang-Jun Park, Yeol Kyo Choi, Taeyong Park, Maham Tanveer, Yiwei Cao, Nathan R. Kern, Jumin Lee, Min Sun Yeom, Tristan I. Croll, Chaok Seok, Wonpil Im

## Abstract

This technical study describes all-atom modeling and simulation of a fully-glycosylated full-length SARS-CoV-2 spike (S) protein in a viral membrane. First, starting from PDB:6VSB and 6VXX, full-length S protein structures were modeled using template-based modeling, de-novo protein structure prediction, and loop modeling techniques in GALAXY modeling suite. Then, using the recently-determined most occupied glycoforms, 22 N-glycans and 1 O-glycan of each monomer were modeled using Glycan Reader & Modeler in CHARMM-GUI. These fully-glycosylated full-length S protein model structures were assessed and further refined against the low-resolution data in their respective experimental maps using ISOLDE. We then used CHARMM-GUI Membrane Builder to place the S proteins in a viral membrane and performed all-atom molecular dynamics simulations. All structures are available in CHARMM-GUI COVID-19 Archive (http://www.charmm-gui.org/docs/archive/covid19), so researchers can use these models to carry out innovative and novel modeling and simulation research for the prevention and treatment of COVID-19.

## INTRODUCTION

The ongoing COVID-19 pandemic is affecting the whole world seriously, with near 5 million reported infections and over 300,000 deaths as of May, 2020. Worldwide efforts are underway towards development of vaccines and drugs to cope with the crisis, but no clear solutions are available yet. The spike (S) protein of SARS-CoV-2, the causative virus of COVID-19, is highly exposed outwards on the viral envelope and plays a key role in pathogen entry. S protein mediates host cell recognition and viral entry by binding to human angiotensin converting enzyme-2 (hACE2) on the surface of the human cell^1^.

As shown in **Figure 1A**, S protein is made up of two subunits (termed S1 and S2) that are cleaved at Arg685-Ser686 by the cellular protease furin^2^. The S1 subunit contains the signal peptide, N terminal domain (NTD), and receptor binding domain (RBD) that binds to hACE2. The S2 subunit comprises the fusion peptide, heptad repeats 1 and 2 (HR1 and HR2), transmembrane domain (TM), and cytoplasmic domain (CP). S protein forms a homo-trimeric complex and is highly glycosylated with 22 predicted N-glycosylated sites and 4 predicted O-glycosylated sites^3-4^ (**Figure 1B**), among which 17 N-glycan sites were confirmed by cryo-EM studies^5-6^. Glycans on the surface of S protein could inhibit recognition of immunogenic epitopes by the host immune system. Steric and chemical properties of viral surface are largely dependent upon glycosylation pattern, making development of vaccines targeting S protein even more difficult.

**Figure 1.**
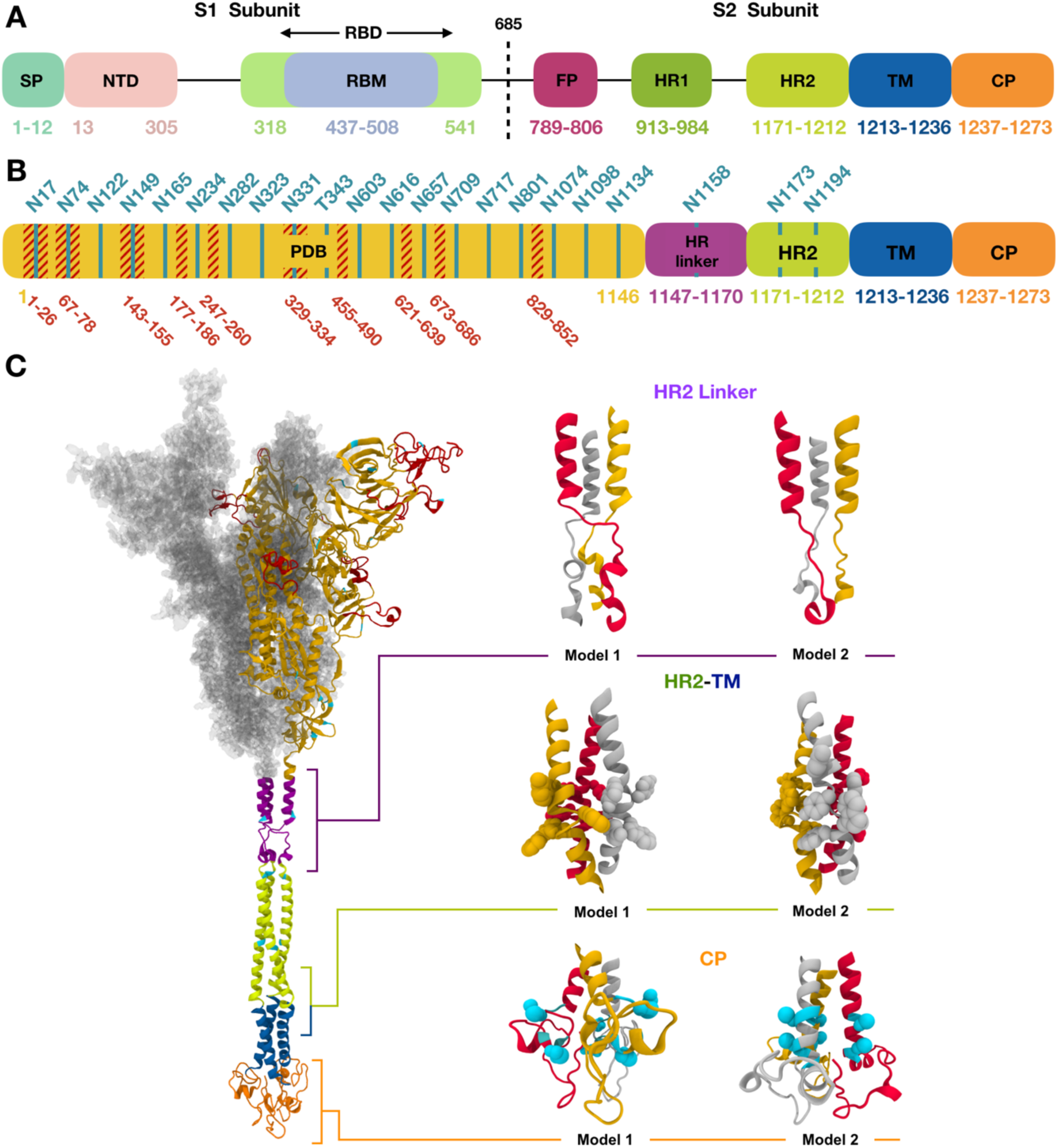
(A) Assignment of functional domains in SARS-CoV-2 S protein: signal peptide (SP), N terminal domain (NTD), receptor binding domain (RBD), receptor binding motif (RBM), fusion peptide (FP), heptad repeat 1 (HR1), heptad repeat 2 (HR2), transmembrane domain (TM), and cytoplasmic domain (CP). (B) Assignment of modeling units used for model building. Glycosylation sites are indicated by residue numbers at the top. Missing loops longer than 10 residues or including a glycosylation site in PDB:6VSB chain A are highlighted in red. Modeled glycosylation sites are shown in cyan. (C) A model structure of full-length SARS-CoV S protein is shown on left panel using the domain-wise coloring scheme in (B). For the PDB region, only one chain is represented by a secondary structure, while the other chains are represented by the surface. Two models are selected for each HR linker, HR2-TM, and CP domain are enlarged on the right panel of (C). Trp and Tyr in HR2-TM are shown in spheres, which are key residues placed on a plane to form interactions with the lipid head group. For CP domain models, the Cys cluster is known to have high probabilities of palmitoylation. Cys1236 and Cys1241 for model 1 and Cys1236 and Cys1240 for model 2 are selected for palmitoylation sites in this study and are represented as cyan spheres. Illustration of S proteins were generated using VMD^14^.

Structures of the RBD complexed with hACE2 have been determined by X-ray crystallography^7-9^ and cryo-EM^10^. Structures corresponding to RBD-up (PDB:6VSB)^5^ and RBD-down (PDB:6VXX)^6^ states of glycosylated S protein were reported by cryo-EM. Molecular simulation studies based on the glycosylated S protein cryo-EM structures have also been reported^11-12^. However, missing domains, residues, disulfide bonds, and glycans in the experimentally resolved structures make it extremely challenging to understand S protein structure and dynamics at the atomic level. For example, 533 residues are missing in PDB:6VSB (**Figure 1B**), and structures of HR2, TM, and CP domains are not available.

In this study, we report all-atom fully-glycosylated, full-length S protein structure models that can be easily used for further molecular modeling and simulation studies. Starting from PDB:6VSB and 6VXX, the structures were generated by combined endeavors of protein structure prediction of missing residues and domains, in silico glycosylation on all potential sites, and refinement based on experimental density maps. In addition, we have built a viral membrane system of the S proteins and performed all-atom molecular dynamics simulation to demonstrate the usability of the models.

All S protein structures and viral membrane systems are available in CHARMM-GUI^13^ COVID-19 Archive (http://www.charmm-gui.org/docs/archive/covid19), so they can serve as a starting point for studies aiming at understanding biophysical properties and molecular mechanisms of the S protein functions and its interactions with other proteins. The computational studies described in this study can also be an effective tool for rapidly coping against possible current and future genetic mutations in the virus.

### FULL-LENGTH SARS-COV-2 S PROTEIN MODEL BUILDING

A schematic view of domain assignment of S protein is provided in **Figure 1A** for functional domains and **Figure 1B** for modeling units. Missing parts in the PDB structures (6VSB and 6VXX) were modeled (colored in red, **Figure 1C**), and structures for four additional modeling units were predicted under the C_3_ symmetry of the homo-trimer. Two structures were selected for each of HR linker, HR2-TM, and CP, resulting in 8 model structures after the domain by domain assembly. Note that the wild-type sequence was used in our models, while 5 and 18 mutations are present in 6VSB and 6VXX, respectively.

First, missing loops in the RBD (residues 336-518) were constructed by template-based modeling using GalaxyTBM^15^. PDB:6M17^10^, which covers the full RBD, was used as a template. Other missing loops in the PDB structures were modeled by FALC (Fragment Assembly and Loop Closure) program^16^ using a light modeling option (i.e., number of generated conformations = 100) except for the loops close to the possible glycosylation sites (residues 67-78, 143-155, 177-186, 247-260, 673-686), for which a heavier modeling option was used with 500 generated conformations. The long N-terminal region (residues 1-26) is not expected to be sampled very well by this method. Some loops and the N-terminus were re-modeled based on the electron density map, as explained below in “Model assessment and refinement”.

Second, an *ab initio* monomer structure prediction and *ab initio* trimer docking were used for the HR linker region (residues 1148-1171 in **Figure 1B**). The single available structural template PDB:5SZS^17^ from SARS-CoV-1 covers only a small portion of the linker, and the resulting template-based structure had a poor trimer interface. Helix and coil regions were first modeled using FALC based on the secondary structure prediction by PSIPRED^18^, and a trimer helix bundle structure was generated by the symmetric docking module of GalaxyTongDock^19^. Two trimer model structures were finally selected after manual inspection (**Figure 1C**).

Third, a template-based modeling method using GalaxyTBM was used to predict the structure of the HR2 domain (residues 1172-1213) using PDB:2FXP^20^ from SARS-CoV-1 as a template. This template has 100% sequence identity and 1.278 similarity with 95.2% sequence coverage based on HHalign^21^.

Fourth, a template-based model was constructed for the TM domain (residues 1214-1237) using GalaxyTBM. PDB:5JYN^22^, a crystal structure of the TM domain of gp-41 protein of HIV, was used as a template. 5JYN has 28% sequence identity and 0.570 sequence similarity with 100% sequence coverage. After the initial model building, manual alignment and FALC loop modeling were applied to locate Trp and Tyr residues on a plane in the final model structure. This was performed to form close interaction of these residues with the lipid head group to be constructed in the membrane building stage (see below). Two models were selected for the HR2-TM junction, one following more closely to the template structure (model 2 in **Figure 1C**) and the other with more structural difference (model 1).

Fifth, the monomer structure of the Cys-rich CP domain (residues 1238-1273) was predicted by GalaxyTBM using PDB:5L5K^23^ as a template. The trimer structure was built by a symmetric *ab initio* docking of monomers using GalaxyTongDock. Loop modeling was performed for the residues missing in the template using FALC. Among the top-scored docked trimer structures, two models with some Cys residues pointing towards the lipid bilayer were selected (**Figure 1C**), considering the possibility of anchoring palmitoylated Cys residues in a lipid bilayer.

Finally, model structures of the above domains were assembled by aligning the C_3_ symmetry axis and modeling domain linkers by FALC. All 8 models for each of 6VSB and 6VXX, generated by assembling each of the two models for three regions (HR2 linker, HR2-TM, and CP), were subject to further refinement by GalaxyRefineComplex^24^ before being attached to the experimentally resolved structure region. The full structure was subject to local optimization by the GALAXY energy function.

### GLYCAN SEQUENCES AND STRUCTURE MODELING

The mass spectrometry data^3-4^ revealed the composition of glycans in each glycosylation site of S protein, and we built glycan structures from their composition. The glycan composition at each site is reported in the form of HexNAc(i)Hex(j)Fuc(k)NeuAc(l), representing the numbers of N-acetylhexosamine (HexNAc), hexose (Hex), deoxyhexose (Fuc), and neuraminic acid (NeuAc). For a given composition, it is possible to generate a large number of glycan sequences with different combinations of residue and linkage types, but this large number can be reduced to a limited number of choices based on the knowledge of biosynthetic pathways of N-linked glycans. All N-glycans share a conserved pentasaccharide core in the form of Manα1-3(Manα1-6)Manβ1-4GlcNAcβ1-4GlcNAcβ, where Man is for D-mannose and GlcNAc is for N-acetyl-D-glucosamine, and they are classified into three types (**Figure 2**): high-mannose, hybrid, and complex, according to the additional glycan residues linked to the core. The glycan sequences selected for 22 N-linked and 1 O-linked glycosylation sites of S protein monomer are shown in **Table 1**. Note that the experiment data show that multiple glycan compositions are detected in every glycosylation site with different abundances^3-4^. For each glycosylation site, we chose the composition with the highest abundance and build the glycan sequence based on the composition. In the case that multiple sequences can be generated from the same composition, we selected the most plausible sequence, as explained in the following examples.

**Table 1.**
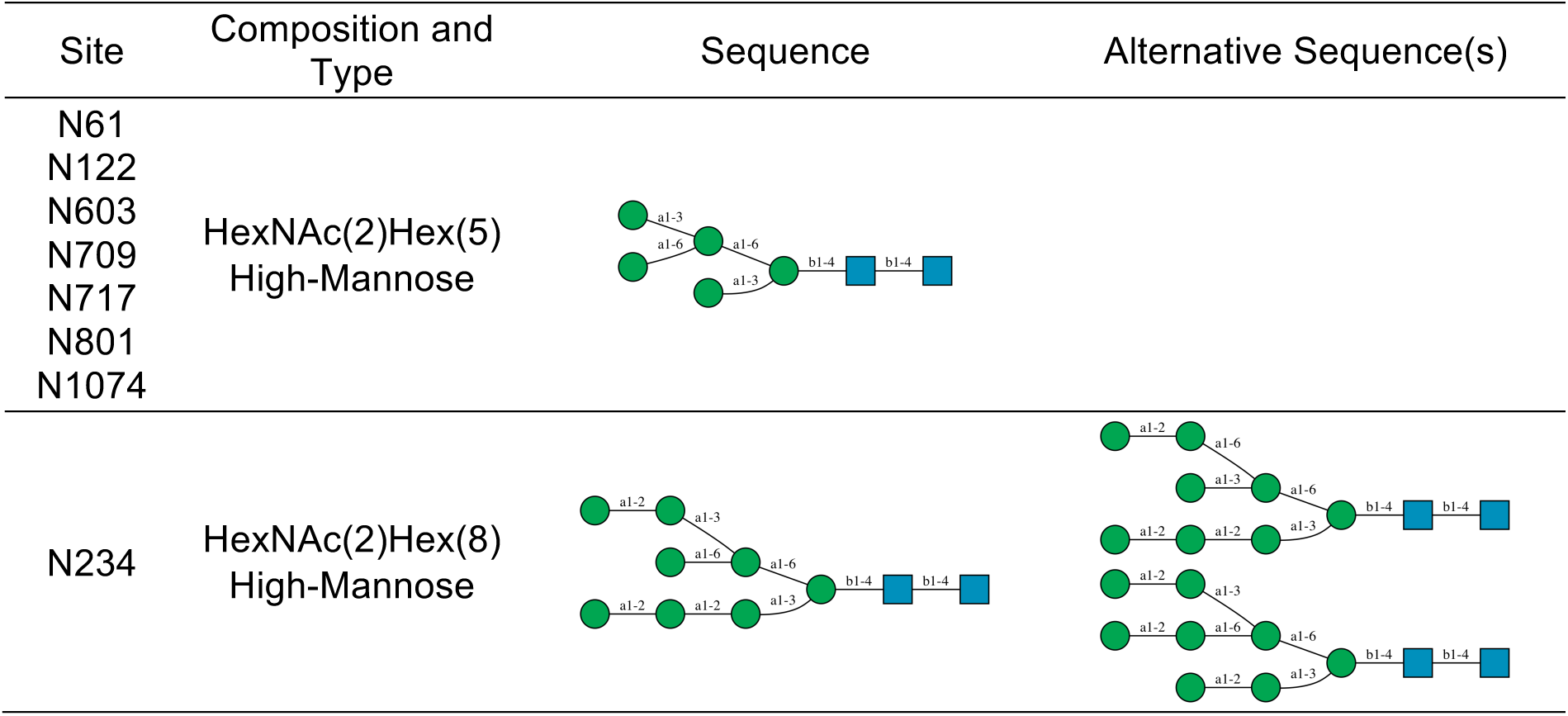
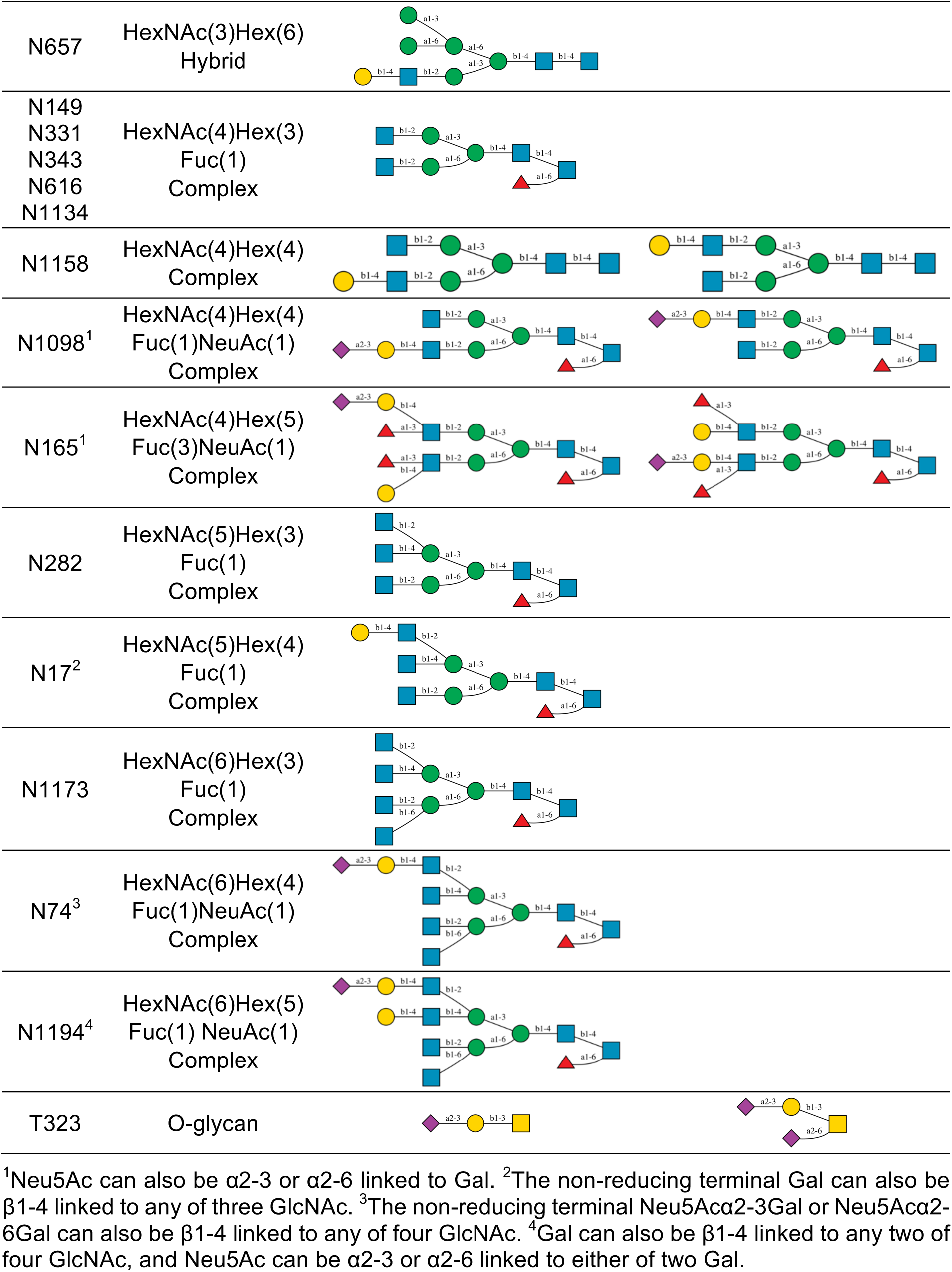
Glycosylation sites and selected glycan compositions and sequences in this study.

| Site | Composition and Type | Sequence | Alternative Sequence(s) |
| --- | --- | --- | --- |
| N61<br>N122<br>N603<br>N709<br>N717<br>N801<br>N1074 | HexNAc(2)Hex(5)<br>High-Mannose |  |  |
| N234 | HexNAc(2)Hex(8)<br>High-Mannose |  |  |

|  |  |
| --- | --- |
| N657 | HexNAc(3)Hex(6)<br>Hybrid |
| N149<br>N331<br>N343<br>N616<br>N1134 | HexNAc(4)Hex(3)<br>Fuc(1)<br>Complex |
| N1158 | HexNAc(4)Hex(4)<br>Complex |
| N1098<sup>1</sup> | HexNAc(4)Hex(4)<br>Fuc(1)NeuAc(1)<br>Complex |
| N165<sup>1</sup> | HexNAc(4)Hex(5)<br>Fuc(3)NeuAc(1)<br>Complex |
| N282 | HexNAc(5)Hex(3)<br>Fuc(1)<br>Complex |
| N17<sup>2</sup> | HexNAc(5)Hex(4)<br>Fuc(1)<br>Complex |
| N1173 | HexNAc(6)Hex(3)<br>Fuc(1)<br>Complex |
| N74<sup>3</sup> | HexNAc(6)Hex(4)<br>Fuc(1)NeuAc(1)<br>Complex |
| N1194<sup>4</sup> | HexNAc(6)Hex(5)<br>Fuc(1) NeuAc(1)<br>Complex |
| T323 | O-glycan |

<sup>1</sup>Neu5Ac can also be α2-3 or α2-6 linked to Gal. <sup>2</sup>The non-reducing terminal Gal can also be β1-4 linked to any of three GlcNAc. <sup>3</sup>The non-reducing terminal Neu5Acα2-3Gal or Neu5Acα2-6Gal can also be β1-4 linked to any of four GlcNAc. <sup>4</sup>Gal can also be β1-4 linked to any two of four GlcNAc, and Neu5Ac can be α2-3 or α2-6 linked to either of two Gal.

**Figure 2.**
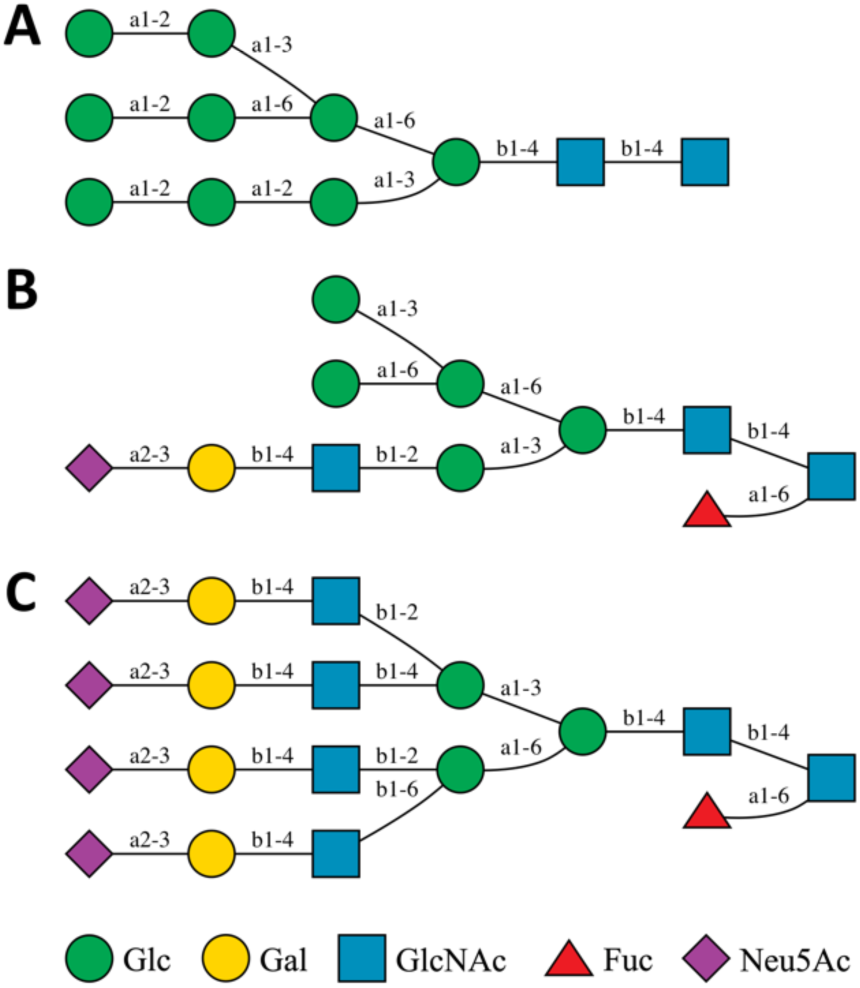
Examples of three types of N-linked glycans: (A) high-mannose, (B) hybrid, (C) complex.

The high-mannose type with composition HexNAc(2)Hex(5) in several glycosylation sites (Asn61, Asn122, Asn603, Asn709, Asn717, Asn801, and Asn1074) have multiple possible sequences. The most common one has two mannose residues linked to the Manα1-6 arm of the core by α1-3 and α1-6 linkages, respectively, but it is also possible that a Manα1-2Man motif is α1-2 linked to the Manα1-3 arm or α1-3/α1-6 linked to the Manα1-6 arm. In this study, the first sequence (**Table 1**) was selected because it is more plausible than others based on the N-glycan synthetic pathway. The complex type with composition HexNAc(4)Hex(4) at Asn1158 has multiple sequences and two of them are equally feasible. Starting from the common core, two additional GlcNAc can be added to the non-reducing terminus. Although it is possible that both GlcNAc are linked to the same arm (α1-3 or α1-6 arm), it is considered to be more likely that each arm has one GlcNAc attached. Then, an additional D-galactose (Gal) can be linked to either α1-3 or α1-6 arm, resulting in two equally possible sequences shown in in **Table 1**. In this study, we selected the sequence with galactose linked to the α1-6 arm for Asn1158 in all chains. However, readers can use Glycan Reader & Modeler^25-27^ in CHARMM-GUI to modify the glycan sequences and build structures according to their requirements.

For the N-glycans containing L-fucose (Fuc) and 5-N-Acetyl-D-neuraminic acid (Neu5Ac), a variety of sequences result from different positions of Fuc and Neu5Ac. Fuc can be α1-6 linked to the reducing terminal GlcNAc or α1-3 linked to GlcNAc in the non-reducing terminal motif Galβ1-4GlcNAc (Asn165 in **Table 1**). We prioritized to attach Fuc to the reducing terminus and considered the non-reducing terminus only if there were more than one Fuc residue. Neu5Ac is attached to Gal in the non-reducing terminal motif Galβ1-4GlcNAc by an α2-3 or α2-6 linkage. In spite of multiple possible sequences, we only chose one of them to be attached to the corresponding glycosylation site. In our model, there is only one O-linked glycan at Thr323, and we selected the sequence Neu5Acα2-3Galβ1-3GalNAcα.

Finally, we used Glycan Reader & Modeler in CHARMM-GUI to model glycan structures at each glycosylation site based on the glycan sequence in **Table 1**. Glycan Reader & Modeler uses a template-based glycan structure modeling method.

### MODEL ASSESSMENT AND REFINEMENT

Ideally, the starting coordinates for any atomistic simulation should closely represent a single “snapshot” from the ensemble of conformations taken on by the real-world structure. For regions where high-resolution experimental structural information is missing, care should be taken to ensure the absence of very high-energy states and/or highly improbable states separated from more likely ones by large energy barriers or extensive conformational rearrangements. Examples of the latter include local phenomena such as erroneous *cis* peptide bonds, flipped chirality, bulky sidechains trapped in incorrect conformations in packed environments, and regions modelled as unfolded/unstructured where in reality a defined structure exists. While some issues may be resolved during equilibrium simulation, in many cases such errors will persist throughout simulations of any currently-accessible timescale, biasing the results in unpredictable ways.

With the above in mind, when preparing an experimentally-derived model for simulation, it is important to remember that the coordinates deposited in the PDB are themselves an imperfect representation of reality. In particular, at relatively low resolutions where the S protein structures were determined, some level of local coordinate error is to be expected^28-29^. While the majority of these tend to be somewhat trivial mistakes that can be fixed by simple equilibration, the possibility of more serious problems cannot be ruled out. While both 6VSB and 6VXX were of unusually high quality for their resolutions (perhaps a testament to the continuous improvement of modelling standards), there were nevertheless some issues that without correction could lead to serious local problems in an equilibrium simulation.

An example of this is the loop-spanning residues 673-686 that are missing in 6VSB (**Figure 3**). In such situations, it is quite common for the “ragged ends” (the modelled residues immediately preceding and following the missing segment) to be quite unreliable. In this case, the three residues C-terminal to the break were incorrectly modelled as a sharp turn to form a tight β-hairpin in 6VSB, leaving a gap compatible only with a single residue where in reality twelve residues were missing. This is likely a failure of automated model-building, as such routines are often confused in situations like this, where strong density at the base of a hairpin is apparently connected, while the true path is weakly resolved due to high mobility. As might be expected, a naïve loop modelling approach based on the as-deposited coordinates was highly problematic at this site, introducing severe clashes and extremely strained geometry in order to contort the chain into a space that was fundamentally too small. Modelling with FALC yielded a loop model with significantly improved geometry and interactions with the local environment, which was then further interactively remodeled into the low-resolution features of the map as described below.

**Figure 3.**
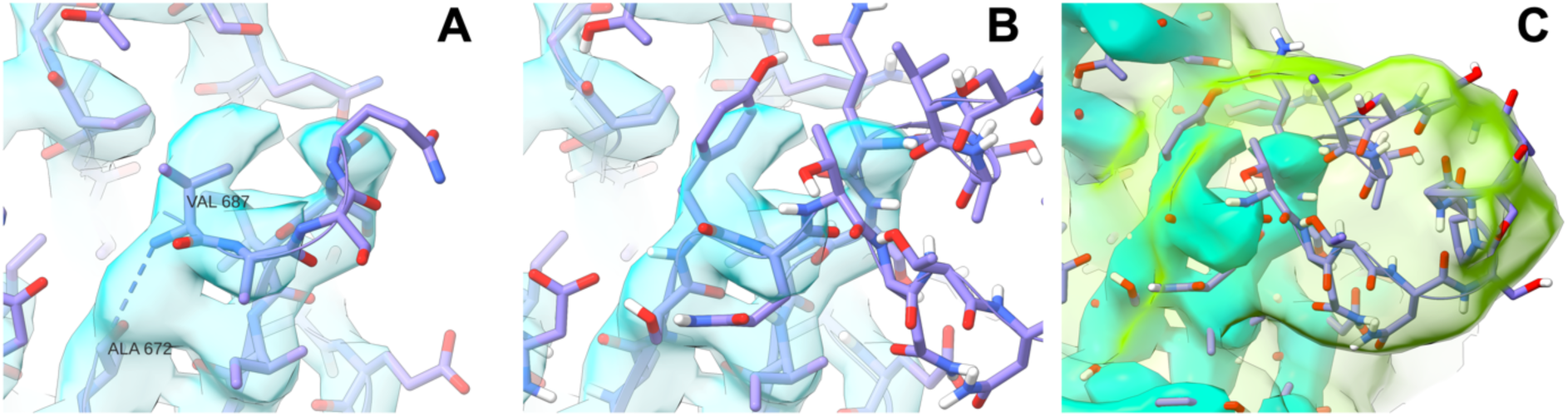
Errors in the experimental model can confound naïve loop modelling approaches. (A) In PDB:6VSB, residues 673-686 were missing, and residues 687-689 were incorrectly modelled forming the appearance of a severely truncated hairpin. Initial attempts at automated modelling of this loop generated numerous clashes and extremely strained geometry. (B) Manual rebuilding into the high-resolution density showed that the space originally modelled as Val687 actually belongs to Tyr674. (C) While the remainder of the loop was not visible in the high-resolution details of the map, the application of strong Gaussian smoothing revealed a clear envelope, suggesting that the hairpin is capped by a short α-helix spanning residues 680-685. The map shown in cyan is as-deposited and contoured at 15σ (all panels), and the green map in (C) is smoothed with a B-factor of 100 Å^2^ and contoured at 5σ. All images were generated using UCSF ChimeraX^30^.

The software package ISOLDE^29^ combines GPU-accelerated interactive molecular dynamics flexible fitting (MDFF) based on OpenMM^31-32^ with real-time validation of common geometric problems, allowing for convenient inspection and correction or triage of prospective models prior to embarking on long simulation. For each of 6VSB and 6VXX, the part of the model covered by the map (protein residues 1-1146) was inspected residue-by-residue in ISOLDE, and was remodeled as necessary to maximize fit of both protein and glycans to the map and resolve minor local issues such as flipped peptide bonds and incorrect rotamers. The applied MDFF potential was a combination of the map as deposited, and a second map smoothed with a B-factor of 100 Å^2^ to emphasize the low-resolution features. The coupling constant of sugar atoms to the MDFF potential was set to one tenth that of the protein atoms to reflect the fact that the majority of sugar residues were highly mobile and effectively unresolved, with typically only 1-3 “stem” residues visible.

For the majority of each model, only minor local changes were necessary, but more extensive work was needed in the sites of *ab initio* loop modelling. These are summarized in **Figure 4** using 6VXX as an example; conformations of these sites modelled in 6VSB were very similar. In general, the starting conformations of the loops modelled *ab initio* were significantly more extended and unstructured than what the map density indicated, suggesting that there may be value in adding a fit-to-density term (where applicable) to the loop modelling algorithm in future.

**Figure 4.**
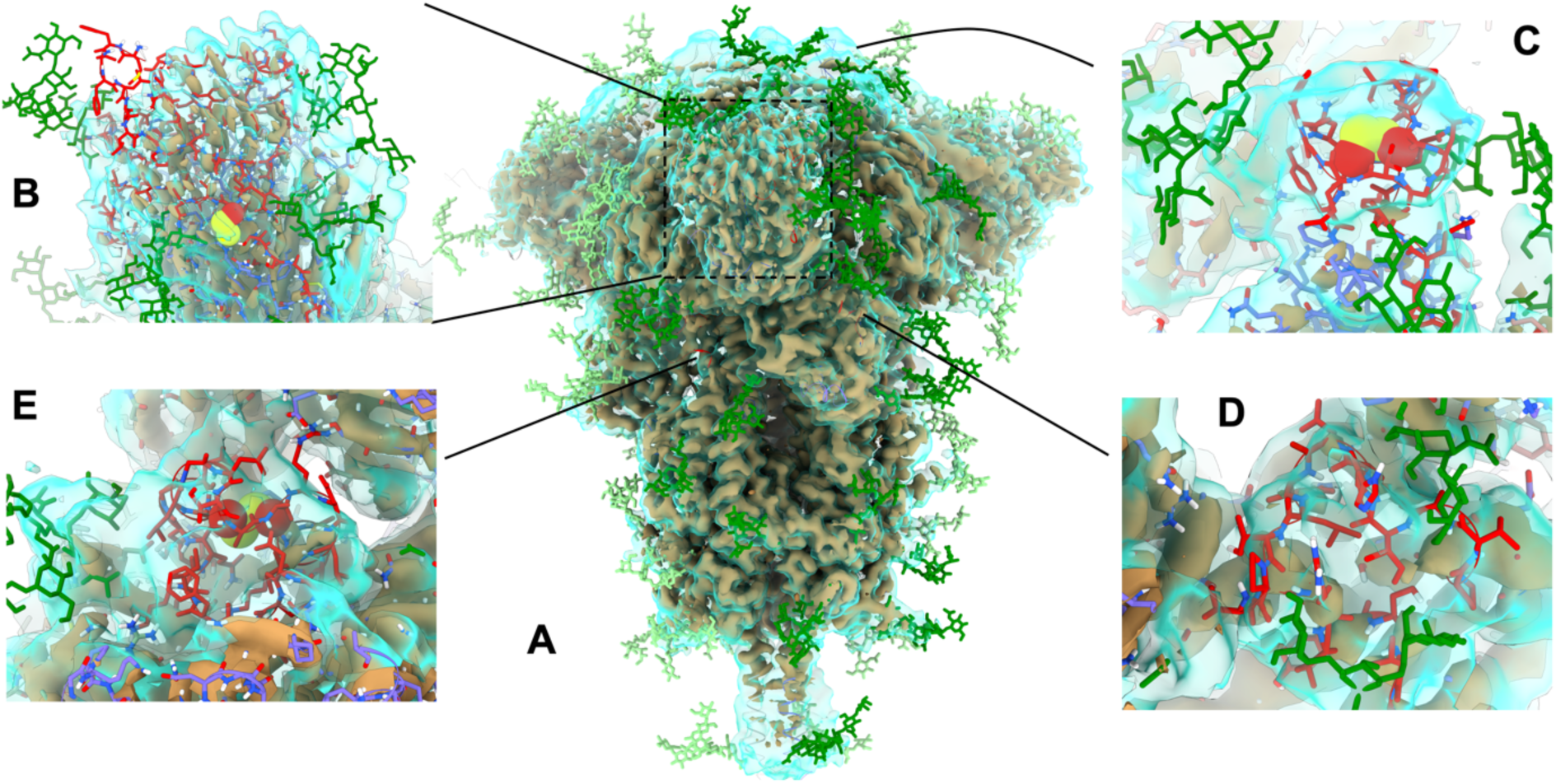
Overview of key sites of extensive remodeling of 6VXX in ISOLDE. In all panels, the tan, opaque surface is the as-deposited map contoured at 9σ, while the transparent cyan surface is a smoothed map (B=100 Å^2^) at 5σ. Glycans are colored in green (dark = chain A; light = chains B and C); newly-added protein carbon atoms in red; pre-existing protein carbons in purple (chain A) or orange (chain B). (A) Overview, highlighting sites of remodeling. (B) The N-terminal domain is exceedingly poorly resolved compared to the remainder of the structure, and the distal face is essentially uninterpretable in isolation. Rebuilding was aided by reference to a crystal structure of the equivalent domain from SARS-CoV-1 (PDB:5X4S^33^), but the result remains the lowest-confidence region of the model. Note in particular the addition of a disulfide bond between Cys15 and Cys136. (C) Extended loop of RBD (residues 468-489). While not visible in the high-resolution data of either cryo-EM map, the conformation seen in an RBD-ACE2 complex (PDB:6M17^10^) was consistent with the low-resolution envelope. (D) Residues 620-641 and (E) 828-854 were associated with strong density indicating a compact structure, but too poorly resolved to support an unambiguous assignment based on the density alone. Using ISOLDE’s interactive MDFF, each loop was packed into a compact fold with reasonable geometry and a hydrophobic core, and suggested with high confidence the presence of a disulfide bond between Cys840 and Cys851. Illustration of S proteins and maps were generated using UCSF ChimeraX.

In this context, it is worth nothing a few issues affecting CoV-2 S protein trajectories recently shared by DE Shaw Research (DESRES)^12^ For example, the starting model for the “closed” trajectory (DESRES-ANTON-11021566) contains 3 or 4 spurious non-proline *cis* peptide bonds in each chain (His245-Arg246, Arg246-Ser247 and Ser640-Asn641 in all chains; Gln853-Lys854 in chains A and C). All but one of these is still present in the final frame of the 10-μs trajectory; His245-Arg246 in chain B flipped to *trans* in the course of simulation. The starting model for the “open” state (DESRES-ANTON-11021571) has *cis* bonds at His245-Arg246, Arg246-Ser247 and Ser640-Asn641 in all chains and Gln853-Lys854 in chain A. At the end of 10-μs, only Ser640-Asn641 and Gln853-Lys854 in chain A were flipped to *trans*. While individually small, local conformation errors like these may be problematic in that they introduce subtle biases throughout the simulation trajectory, a spurious *cis* peptide bond, for example, introduces an unnatural kink in the protein backbone, effectively preventing the formation of any regular secondary structure in the immediate vicinity.

The Gln853-Lys854 bond is found at the C-terminal end of the 828-854 loop, which is missing in all current experimental structures. While the local resolution of the cryo-EM maps in this region is low, the density associated with chain A of 6VSB in particular strongly suggests residues 849-856 are α-helical, and that Cys851 forms a disulfide bond to Cys840 (**Figure 5A**).

**Figure 5.**
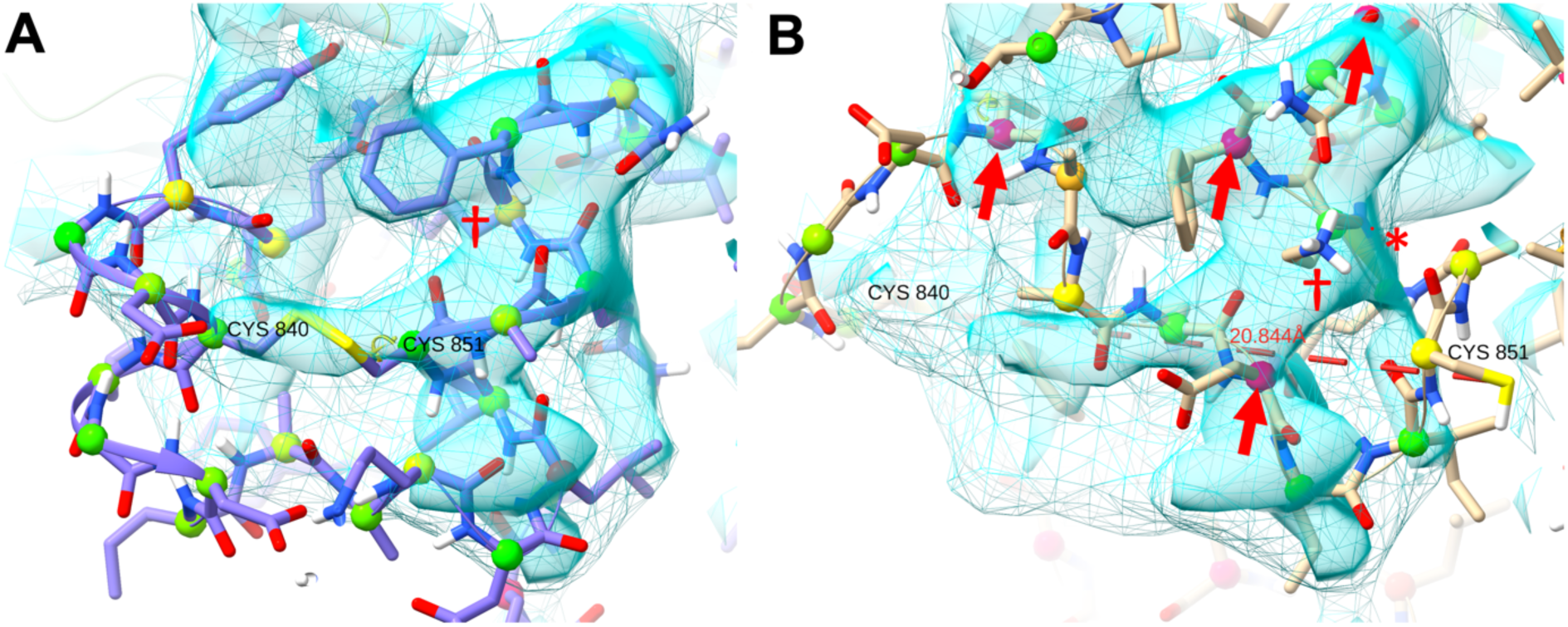
In chain A of PDB:6VSB, the density strongly suggests an α-helix spanning residues 849-856, and a disulfide bond between Cys840 and Cys851. (A) As modelled in this study, colored balls on CA atoms are ISOLDE markup indicating Ramachandran status (green = favored; hot pink = outlier). Lysine 854 (dagger) is pointing into the page, where it forms a salt bridge with Asp614 of an adjacent chain. (B) The starting chain A model of DESRES-ANTON-11021571 has Cys840 and Cys851 about 20 Å apart. There are four severe Ramachandran outliers (red arrows), and a *cis* peptide bond at 853-854 (asterisk). Lysine 854 points outwards through the core of the helical density, and is blocked from the position in our model by the steric bulk of Phe855. All figures were generated using UCSF ChimeraX

We therefore modeled it as such for all chains of all models. Lys854 appears to play a structural role, salt-bridging to Asp614 of the adjacent chain and further H-bonding to backbone oxygens within its own chain. In the starting model of DESRES-ANTON-11021571 (**Figure 5B**), this residue is instead solvent-exposed with its sidechain facing in the opposite direction, distorting the backbone into random coil (including the aforementioned 853-854 *cis* bond) and effectively blocked from returning to the conformation suggested by the density due to the steric bulk of the Phe855 sidechain. Meanwhile, Cys840 and Cys851 are far from each other and blocked from approach by many intervening residues. While residues 849-856 did settle to a helical conformation in chain A (but not chains B and C) during the DESRES simulation, the cysteine sulfur atoms remained 13-16 Å apart at the end of the simulation.

### COVID-19 ARCHIVE IN CHARMM-GUI (http://www.charmm-gui.org/docs/archive/covid19)

All S protein models and simulation systems are available in this archive. The model name follows the model numbers used for HR linker, HR2-TM, and CP structures (**Figure 1C**). For example, “6VSB_1_1_1” represents a model based on PDB:6VSB with HR linker model 1, HR2-TM model 1, and CP model 1. Each PDB file contains all glycan models and their CONECT information for its reuse in CHARMM-GUI. Also, note that all disulfide bond information is included in each PDB file. Therefore, both glycans and disulfide bonds are automatically recognized when the PDB file is uploaded to CHARMM-GUI PDB Reader^34^. For simulation systems (see the next section for details), we provide the simulation system and input files for CHARMM^35^, NAMD^36^, GROMACS^37^, GENESIS^38^, AMBER^39^, and OpenMM^31-32^.

### MEMBRANE SYSTEM BUILDING

*Membrane Builder*^40-44^ in CHARMM-GUI was used to build a viral membrane system of a fully-glycosylated full-length S protein with palmitoylation at Cys1236 and Cys1241 for CP model 1 and Cys1236 and Cys1240 for CP model 2 as these residues are solvent-exposed. **Table 2** shows approximate time of each step during the building process. An online video demo explaining the building process in detail is available (http://www.charmm-gui.org/demo/cov-2-s/1).

**Table 2.** Times to complete a viral membrane system building of a fully-glycosylated full-length S protein with palmitoylation at Csy1236 and Cys1241 for 6VSB_1_1_1.

| STEP | Time (hours) |
| --- | --- |
| 1. PDB reading and manipulation | 0.2 |
| 2. Transmembrane orientation | 0.1 |
| 3. System size and packing | 1.1 |
| 4. Membrane, ion, and water box generation | 1.2 |
| 5. Assembly / input generation | 1.8 / 21.5 |

STEP 1 is to read and manipulate a biomolecule from a PDB file. Since our S protein model PDB files contain all glycan structures and their CONECT information, *PDB Reader & Manipulator*^34^ (via *Glycan Reader & Modeler*) recognizes all the modeled glycans and their linkages. One can change the starting and/or ending residue number to model certain soluble domains only for certain chains. In particular, during the manipulation step, one can use different glycoforms at selected glycosylation sites (**Table 1**); watch video demos for *Glycan Reader & Modeler*. Palmitoylation at certain Cys residues can be made by choosing CYSP (a palmitoylated Cys residue in the CHARMM force field) under “Add Lipid-tail” in this step. In this study, among many palmitoylation options in the S protein CP domain^45^, palmitoylation was made at Cys1236 and Cys1241 for CP model 1 (**Figure 6A**) and Cys1236 and Cys1240 for CP model 2. Because S protein is a transmembrane protein and we want the palmitoyl group to be in the membrane hydrophobic core, the “is this a membrane protein?” option needs to be turned on. Since the CYSP residue does not exist in the Amber force fields, the palmitoylation option should not be used if someone intends to do Amber simulation with the Amber force fields. In any case, it is important to save JOB ID, so that one can check the job progress using *Job Retriever*.

**Figure 6.**
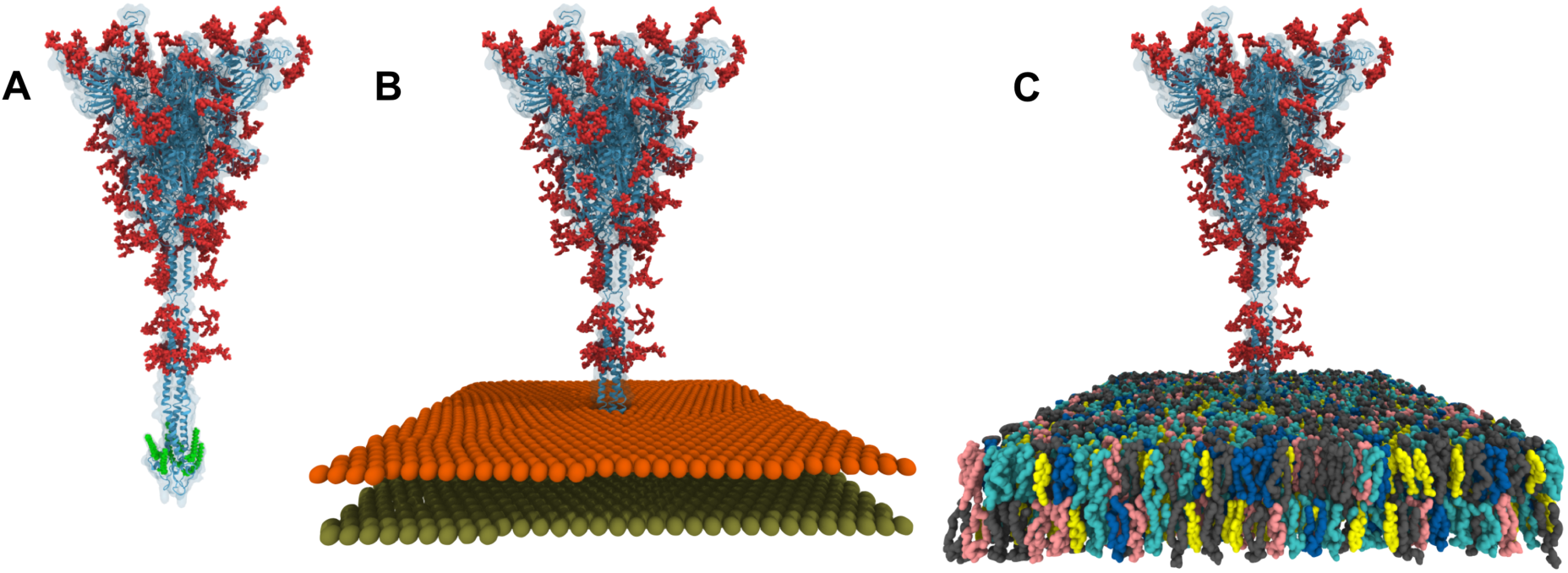
Molecular graphics of 6VSB_1_1_1 system structures after (A) STEP 1, (B) STEP 3, and (C) STEP 5 in Membrane Builder. For clarity, water molecules and ions are not shown in (C). All figures were generated using VMD.

STEP 2 is to orient the TM domain into a bilayer. By definition, a bilayer has its normal along the Z axis and its center at Z = 0. The TM domain of each S protein model was pre-oriented by aligning the TM domain’s principal axis along the Z axis and its center at Z =0 and by positioning the S1 domain at Z > 0. Therefore, there is no need to change the orientation; however, one can translate S protein along the Z axis for different TM positioning. One should always visualize “step2_orient.pdb” to make sure that the protein is properly positioned in a bilayer.

To build any homogeneous or heterogeneous bilayer system, *Membrane Builder* uses the so-called “replacement” method that first packs lipid-like pseudo atoms (STEP 3), and then replaces them with lipid molecules one at a time by randomly selecting a lipid molecule from a lipid structural library (STEP 4). In this study, the system size along the X and Y axes was set to 250 Å to have enough space between S1/S2 domains in the primary system and its image system. The system size along the Z axis was determined by adding 22.5 Å to the top and bottom of the protein, yielding an initial system size of ∼250 × 250 x 380 Å^3^. Although biological viral membranes are asymmetric with different ratios of lipid types in the inner and outer leaflets^46-47^, in this study, the same mixed lipid ratio (DPPC:POPC:DPPE:POPE:DPPS:POPS:PSM:Chol=4:6:12:18:4:6:20:30) was used in both leaflets to represent a liquid-ordered viral membrane based on lipidomic data from influenza and HIV^46-47^; see **Table 3** for the full name of each lipid and their number in the system with the CP model 1. One should always visualize “step3_packing.pdb” (**Figure 6B**) to make sure that the protein is nicely packed by the lipid-like pseudo atoms and the system size in XY is reasonable.

**Table 3.** Number of lipids in the S protein membrane system.

| Lipid type<sup>1</sup> | Outer leaflet<sup>2</sup> | Inner leaflet<sup>2</sup> |
| --- | --- | --- |
| DPPC | 48 | 46 |
| POPC | 72 | 69 |
| DPPE | 144 | 138 |
| POPE | 216 | 207 |
| DPPS | 48 | 46 |
| POPS | 72 | 69 |
| PSM | 240 | 230 |
| CHOL | 360 | 345 |

<sup>1</sup>DPPC for di-palmitoyl-phosphatidyl-choline; POPC for palmitoyl-oleoyl-phosphatidyl-choline; DPPE for di-palmitoyl-phosphatidyl-ethanolamine; POPE for palmitoyl-oleoyl-phosphatidyl-ethanolamine; DPPS for di-palmitoyl-phosphatidyl-serine; POPS for palmitoyl-oleoyl-phosphatidyl-serine; PSM for palmitoyl-sphingomyelin; CHOL for cholesterol. <sup>2</sup>The numbers of lipids in the upper and lower leaflets are different due to the area of the CP domain (model 1) embedded in the lower leaflet. For the CP model 2, the lower leaflet has the same number of lipids as in the upper leaflet.

For a large liquid-ordered system (with high cholesterol) like the current system, the replacement in STEP 4 takes long time due to careful checks of lipid bilayer building to avoid unphysical structures, such as penetration of acyl chains into ring structures in sterols, aromatic residues, and carbohydrates. This is the reason why water box generation and ion placement are done separately in STEP 4.

STEP 5 consists of assembly and input generation. Water molecules overlapping with S protein, glycans, and lipids were removed during the assembly (**Figure 6C**). While we chose GROMACS together with the CHARMM force field^48-51^ for this study, one can choose other simulation programs or use the Amber force fields^52-56^ for Amber simulation. Note that input generation takes long time to prepare all restraints for protein positions, chiral centers, cis double bonds of acyl tails, and sugar chair conformations; these restraints are used for the optimized equilibration inputs to prevent any unwanted structural changes in lipids and embedded proteins, so that production simulations can probe physically realistic behaviors of biomolecules.

### All-Atom MD SIMULATION

In this study, we used the CHARMM force field for proteins, lipids, and carbohydrates. TIP3P water model^52, 57^ was used along with 0.15 M KCl solution. The total number of atoms is 2,343,394 (6VSB_1_1_1: 668,899 water molecules, 2,128 K^+^, and 1,857 Cl^-^); see COVID-19 archive to see other system information. The van der Waals interactions were smoothly switched off over 10–12 Å by a force-based switching function^58^ and the long-range electrostatic interactions were calculated by the particle-mesh Ewald method^59^ with a mesh size of ∼1 Å.

We used GROMACS 2018.6 for both equilibration and production with LINCS^60^ algorithm using the inputs provided by CHARMM-GUI^43^. To maintain the temperature (310.15 K), a Nosé-Hoover temperature coupling method^61-62^ with a tau-t of 1 ps was used, and for pressure coupling (1 bar), a semi-isotropic Parrinello-Rahman method^63-64^ with a tau-p of 5 ps and a compressibility of 4.5 × 10^−5^ bar^-1^ was used. During equilibration run, NVT (constant particle number, volume, and temperature) dynamics was first applied with a 1-fs time step for 250 ps. Subsequently, the NPT (constant particle number, pressure, and temperature) ensemble was applied with a 1-fs time step (for 2 ns) and with a 2-fs time step (for 18 ns); note that this is the only change that we made from the input files generated by CHARMM-GUI for longer equilibration due to the system size. During the equilibration, various positional and dihedral restraint potentials were applied and their force constants were gradually reduced. No restraint potential was used for production.

All equilibration steps were successfully completed for all 16 systems. We were able to perform a production run for 6VSB_1_1_1 with a 4-fs time-step using the hydrogen mass repartitioning technique^65^. A 30-ns production movie shown in **Movie S1** demonstrates that our models and simulation systems are suitable for all-atom MD simulation. The results of these simulations will be published elsewhere.

## CONCLUDING DISCUSSION

All-atom modeling of a fully-glycosylated full-length SARS-CoV-2 S protein structure turned out to be a challenging project (at least to us) as it required many techniques such as template-based modeling, de-novo protein structure prediction, loop modeling, and glycan structure modeling. To our surprise, even after initial modeling efforts that were much more than enough for typical starting structure generation for MD simulation, considerable further refinement was necessary to make a reasonable model that was suitable for simulation. While we have done our best, it is possible that our models may still contains some subtle or larger errors / mistakes that we could not catch. In particular, strong conclusions regarding the N-terminal domain should be avoided until a high-resolution experimental structure becomes available to confirm and/or correct the starting coordinates. Nonetheless, we believe that our models are at least good enough to be used for further modeling and simulation studies. We will improve our models as we hear issues from other researchers. It is our hope that our models can be useful for other researchers and development of vaccines and antiviral drugs.

## ACKNOWLEDGMENT

This study was supported in part by grants from NIH GM126140, NSF DBI-1145987 and MCB-1810695, a Friedrich Wilhelm Bessel Research Award from the Humboldt Foundation (WI), National Research Foundation of Korea grants (2019M3E5D4066898 and 2016M3C4A7952630) funded by the Korea government (CS), Wellcome Trust grant number 209407/Z/17/Z (TC), and the National Supercomputing Center with supercomputing resources including technical support (KSC-2020-CRE-0089) (MSY).

## SUPPORTING INFORMATION

In the supplementary material, Movie S1 shows a 30-ns production trajectory of system 6VSB_1_1_1.

## NOTES

The authors declare no competing financial interest.

## TOC Graphics

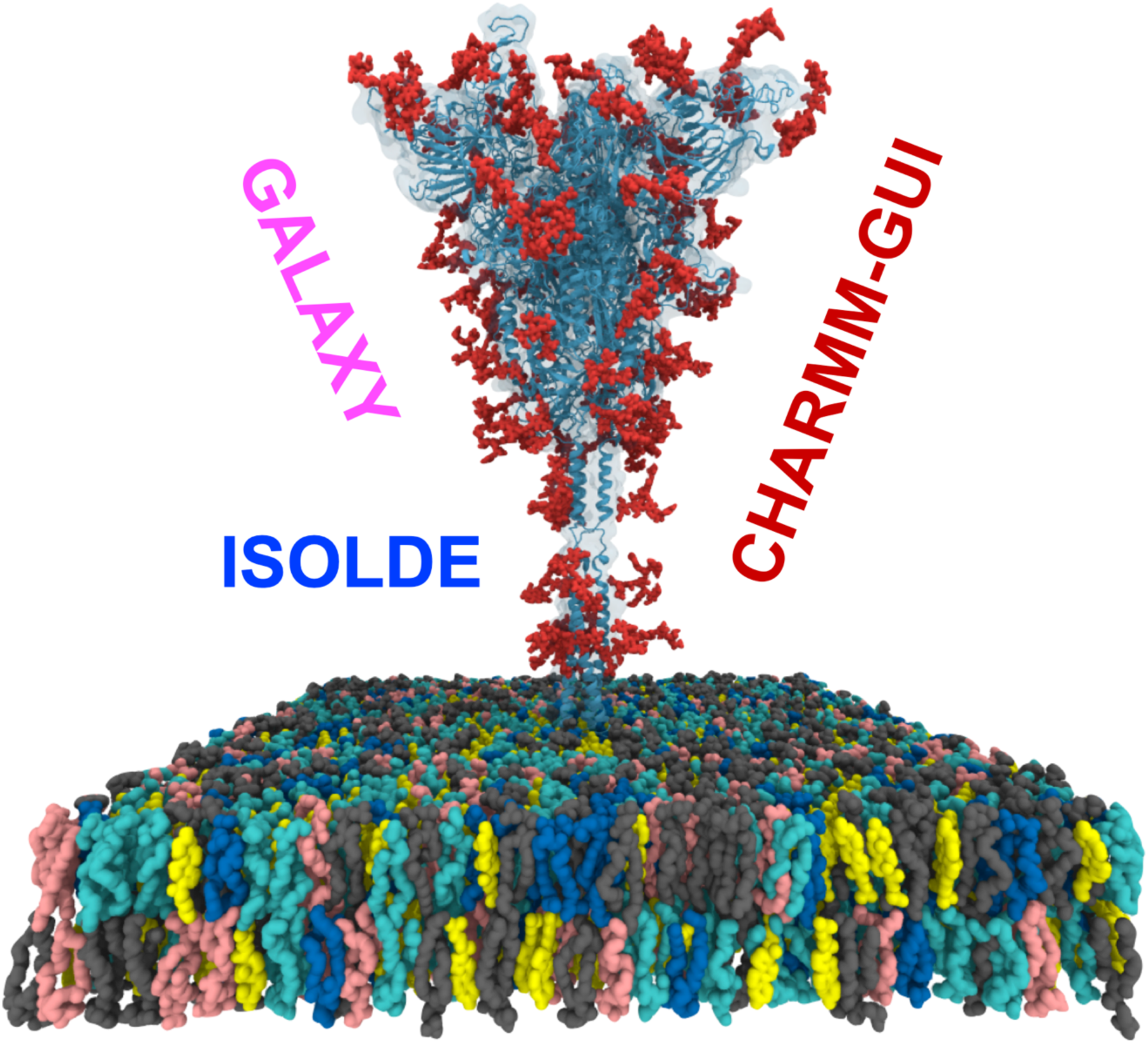

